# Developmental and cell-specific expression of *Cacna1d* splice variants

**DOI:** 10.1101/598722

**Authors:** LaCarubba Brianna, Bunda Alexandra, Savage Kitty, Sargent Hannah, Akiki Marie, Foxall Thomas, Andrade Arturo

**Author notes:** Arturo.Andrade (Corresponding author).

## Abstract

Ca_V_1.3 is an L-type voltage-gated calcium channel implicated in several functions including gene expression, pacemaking activity, and neurotransmitter release. The gene that encodes the Ca_V_α_1_-pore forming subunit of Ca_V_1.3 (*Cacna1d*) is a multi-exon gene that undergoes extensive alternative splicing, which provides functional versatility to this gene across tissues and cell-types. The function and expression of several *Cacna1d* splice variants within the C-terminus have been previously characterized. These splice variants differ in their voltage-dependence of activation, Ca^2+^-dependent inactivation, and their sensitivity to dihydropyridines. However, less is known about alternatively spliced exons in *Cacna1d* located downstream of domain I and upstream of the C-terminus (e11, e22a/e22, e31a/e31b/e32). Here, we performed a systematic study to determine the developmental and cell-specific expression of several *Cacna1d* splice variants. We found that the cassette e11 is upregulated during brain development, and in adult cortical tissue is more abundant in excitatory neurons relative to inhibitory interneurons. This exon is also upregulated upon nerve growth factor (NGF) induced differentiation of pheochromocytoma cells, PC12. At the functional level, the splice variants resulting from e11 alternative splicing (+e11-*Cacna1d* and Δe11-*Cacna1d*) form functional Ca_V_1.3 channels with similar biophysical properties in expression mammalian systems. Of the pair of mutually exclusive exons, e22a and e22, the later dominates at all stages. However, we observed a slight upregulation of e22 from embryonic to adult human brain. A second pair of mutually exclusive exons, e31a and e31b, was also studied. We found that e31a increases during brain development. Finally, the cassette exon 32 is repressed in adult brain tissue.

## 1. INTRODUCTION

More than 90 % of multi-exon genes are subject to alternative splicing in eukaryotic cells (Kelemen et al., 2013; Baralle and Giudice, 2017; Gallego-Paez et al., 2017). This selective inclusion and exclusion of exons abounds in the mammalian nervous system (Porter et al., 2017), and is under the control of RNA binding proteins, or splicing factors (Vuong et al., 2016; Zhang et al., 2016). Expression and activity of splicing factors changes throughout development and across cell-types. As a result, the expression of splice variants is substantially different between newborns and adults, as well as among cellular sub-populations (Calarco et al., 2009; Grabowski, 2011; Fogel et al., 2012; Raj et al., 2014).

The genes that encode the Ca_V_α_1_ pore-forming subunit of voltage-gated calcium channels (*Cacna1*) are multi-exon genes subject to alternative splicing (Liao et al., 2005; Catterall, 2011; Lipscombe et al., 2013; Lipscombe and Andrade, 2015). Ca_V_1.3 channels regulates gene expression, sinoatrial node firing (Mangoni et al., 2003; Torrente et al., 2016), transmitter release from auditory hair cells (Brandt et al., 2003; Baig et al., 2011), and hormone release (Scholze et al., 2001; Marcantoni et al., 2007; Sandoval et al., 2017). Ca_V_1.3 channels have been implicated in various pathologies including Parkinson’s disease, primary aldosteronism and autism spectrum disorder (Chan et al., 2010; Baig et al., 2011; Kang et al., 2012; Reinbothe et al., 2013; Scholl et al., 2013; Pinggera et al., 2015). Interestingly, *Cacna1d* splice variants add functional versatility of Ca_V_1.3 channels across tissues and cell-types.

Ca_V_1.3, like all Ca_V_α_1_ subunits, is organized into four homologous domains (DI-IV). Each domain contains six transmembrane-spanning segments (S1-S6), with a re-entrant loop between S5 and S6, which contains glutamates that are key for the selectivity filter. The amino and carboxyl termini, as well as linker sequences between the DI-II, DII-III, and DIII-IV are cytosolic (Fig. 1A). Sites of alternative splicing of sequences encoding for the N-terminus, DI, DI-II linker, DIII, DIV and C-terminus are present in *Cacna1d* pre-mRNAs. Alternative splicing of sequences encoding for the C-terminus in Ca_V_1.3 have been well characterized, and results in splice isoforms with differences in calcium-dependent inactivation, trafficking, protein-protein interactions, and sensitivity to dihydropyridines (DHPs) (Shen et al., 2006; Singh et al., 2008; Tan et al., 2011; Scharinger et al., 2015; Stanika et al., 2016). However, less is known about the function and expression patterns of several other splice variants derived from the *Cacna1d* pre-mRNA. Here, we determined the developmental and cell-specific expression for *Cacna1d* splice variants that differ in regions located downstream of DI and upstream of the C-terminus. We determined the temporal and cell-specific expression of splice variants encoding for sequences within the DI-II linker (cassette e11) of Ca_V_1.3 in brain. We found that splice variants containing e11 (+e11*-Cacna1d*) are dominant in adult brain, excitatory adult cortical projection neurons (PNs), and PC12 cells differentiated with NGF, whereas splice variants lacking e11 (Δe11-*Cacna1d*) prevail at earlier stages of brain development, inhibitory adult cortical interneurons (INs), and undifferentiated PC12 cells. Functionally, both splice variants generate Ca_V_1.3 channels with similar biophysical properties in expression mammalian systems. We determined the temporal expression of splice variants encoding for sequences within DIII (mutually exclusive exons 22a and 22) and DIV (mutually exclusive exons 31a and 31b, and cassette exon 32) of Ca_V_1.3 in brain tissue. Exon 22a is slightly upregulated in adult human brain. Similarly, e31a increases during brain development. In contrast, splice variants with a fusion of exon 31a and 31b, which are predicted to introduce a premature stop codon, are repressed shortly after birth. Finally, splice variants including the cassette exon 32 are dominant in adult brain tissue.

**Figure 1.**
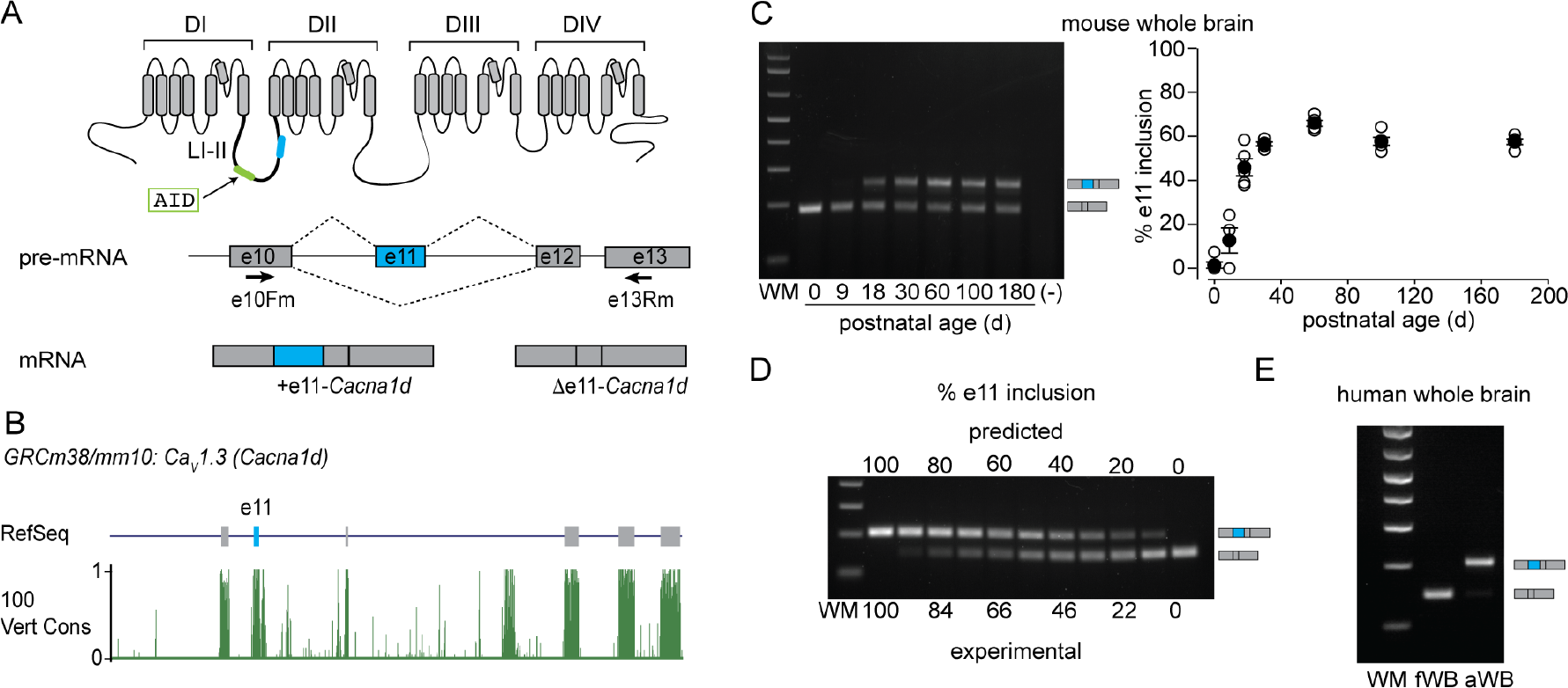
Expression of e11 during postnatal brain development. **A**) *Upper panel:* Schematic of Ca_V_α_1_ pore-forming subunit of voltage-gated calcium channels (DI-IV = domains, LI-II = cytoplasmic linker between DI and DII). Highlighted in light green is the putative location of the alpha-interaction domain (AID), in blue is the putative location of e11. *Lower panel:* Schematic representation of cassette e11 alternative splicing, two splice variants are possible +e11-*Cacna1d* and +Δ11-*Cacna1d*. Boxes represent exons, black lines represent intronic regions, and black arrows represent the approximate location of PCR primers used to amplify +e11-*Cacna1d* and +Δ11-*Cacna1d* splice variants in human and mouse brains. **B**) Visualization of RefSeq Genes and 100 vertebrates Basewise Conservation by 810 PhastCons tracks using UCSC Genome Browser. A region of *Cacna1d* (chr14:30,124,696-30,136,584) from the GRCm38/mm10 mouse assembly is represented. This region contains exons 10, 11, 12, 13, 14, and 15 of the mouse *Cacna1d* gene. In the RefSeq track, vertical gray blocks depict exons and horizontal lines introns. Putative location of e11 in the pre-mRNA is shown with a blue vertical bar. The conservation track displays the PhastCon scores from 0 to 1. **C**) *Left panel:* Representative RT-PCR gel of whole mouse brain at different postnatal ages. Mouse specific primers to amplify both +e11-*Cacna1d* and Δe11-*Cacna1d* were used. *Right panel:* RT-PCRs from male mouse brains (5 per age group). Inclusion of e11 at the following ages are represented as mean (black circles) ± s.e.m. % e11 inclusion. P0 = 2.5 ± 2.5, P9 = 18.4 ± 5.8, P18 = 51 ± 5.4, P30 = 56.1 ± 1.5, P60 = 64.6 ± 1.4, P100 = 58 ± 1.0, P180 = 57 ± 2.3. Empty circles represent individual values. **D**) *Left panel*. Calibration curve to validate quantification of relative amounts ofΔe11 and +e11 by RT-PCR. (−) indicates negative control for PCR (water). **E**) Representative agarose gel of RT-PCR fWB and aWB.

## 2. MATERIALS AND METHODS

### 2.1 Animals and genotyping

All procedures involving vertebrate animals were performed according to Institutional Animal Care and Use Committee protocol at the University of New Hampshire, which follows the National Institutes of Health guide for the care and use of Laboratory animals (NIH Publications No. 8023, revised 1978). Wilde-type C57BL6/J male mice were used in experiments unless indicated otherwise. To label PNs with the red fluorescent protein, tdTomato (tdT), we crossed *CaMKIIα-Cre* mice (*B6.Cg-Tg(Camk2a-cre)T29-1Stl/J*, Jax: 005359) in a mixed C57BL/6 background with *Ai14* mice (*B6.Cg-Gt(ROSA)26Sor*^*tm14(CAG-tdTomato)Hze1*^*/J*, Jax: 007914) in C57BL/6. To label INs with tdT, *Gad2-Cre* mice (*Gad2*^*tm2(cre)Zjh*^/J, Jax: 010802) in C57BL/6 background were crossed with *Ai14* mice. The resulting dual transgenic mouse lines *CaMKIIα;tdT and Gad2;tdT* were heterozygous for both alleles. Conventional toe biopsy was performed on P7-P9 pups. Genomic DNA was extracted with Phire Animal Tissue Direct kit II (ThermoFisher Scientific, F140WH) according to manufacturer instructions. PCR for genotyping was performed with AmpliTaq Gold^®^ 360 mastermix (Thermo Fisher Scientific) using the following conditions: hot start of 95° C for 10 min, followed by 35 cycles (95° C, 30 s; 60° C, 30 s; 72° C, 1 min), and final step of 72° C for 7 min. Primers and expected products are shown in Table 1.

**Table 1.**
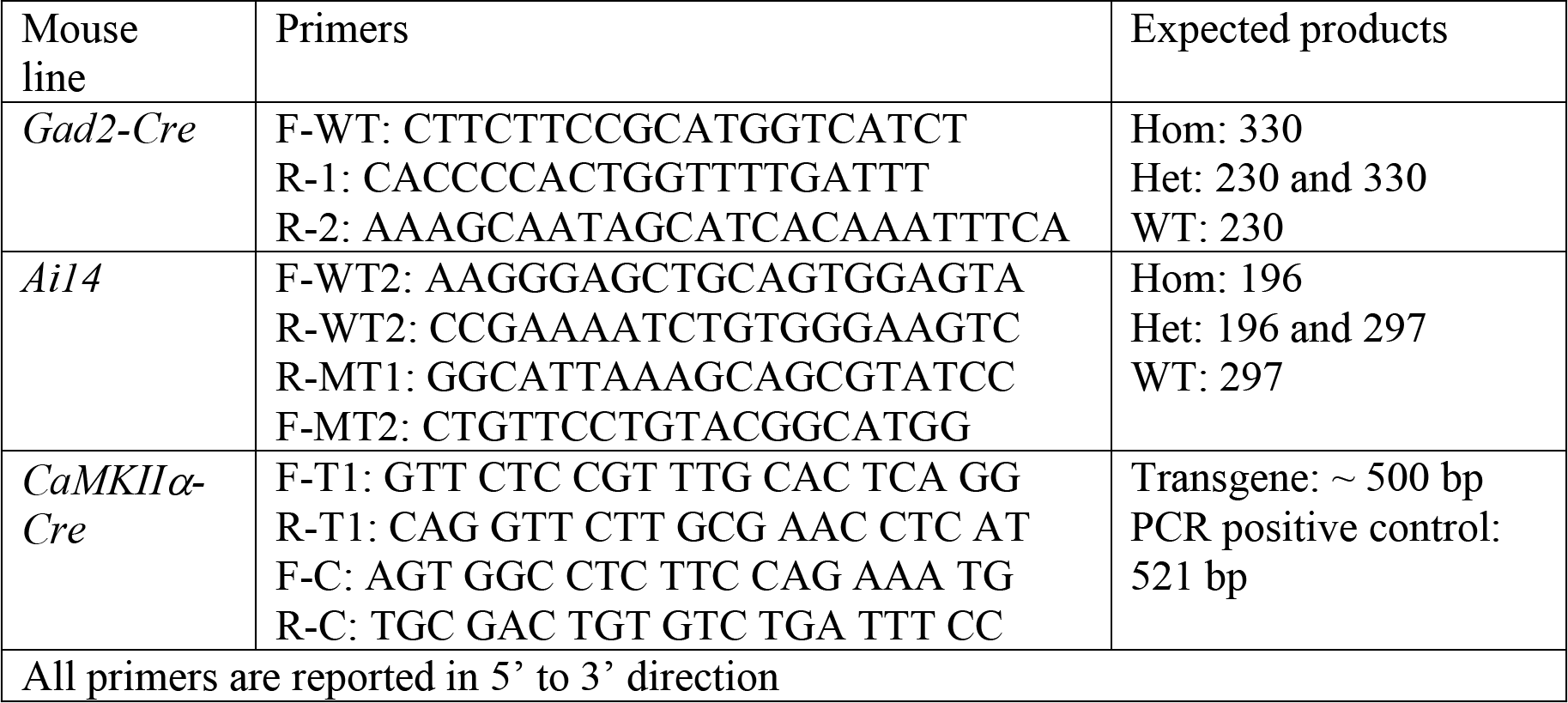
Primers and PCR products for each mouse line.

### 2.2 Sex determination of newborns

To determine the sex of newborn mice, we performed PCR on genomic DNA extracted from tail biopsies. We identified the Y chromosome-specific gene (*Sry*) using the following primers: F, 5'-TGGGACTGGTGACAATTGTC-3, and R, 5'-GAGTACAGGTGTGCAGCTCT-3. As an internal PCR control, we amplified the interleukin-3 (IL-3) gene using the following primers: F, 5'-GGGACTCCAAGCTTCAATCA-3, and R, 5'-TGGAGGAGGAAGAAAAGCAA-3 (Lambert et al., 2000). All primers were added to the same PCR reaction. PCR conditions are as follows: hot start at 95°C for 10 min; followed by 35 cycles of denaturing at 95°C for 30 s, annealing at 56°C for 30 s, and amplification at 72°C for 30s; with a final extension cycle at 72°C for 7 min. Expected size for *Sry* is 402 bp and IL-3 is 544 bp.

### 2.3 Cell culture

Certified PC12 cells (ATCC, CRL1721) were grown in suspension in RPMI-1640 (Gibco, 11875-093) containing 10 % heat-inactivated horse serum (HIHS) (Gibco, 26050088), 5 % fetal bovine serum (FBS) (Gibco, 16000044) and 1 % penicillin/streptomycin (Gibco, 10378016). Cells were passaged twice per week in a 1:2 split. For all our experiments, only cells with 3-15 passages were used. For differentiation experiments, cells were plated on collagen I coated plates (VWR, 62405-603) to promote cell attachment. After 24 hours, growth medium was removed and replaced with low serum differentiation medium, which consisted of RPMI-1640 with 1 % HIHS and 100 ng/mL NGF 2.5S (Alomone, N-100). Medium was replaced every 48 hours for 7 days. After this period, RNA was extracted.

### 2.4 RNA samples

*PC12 cells*. Total RNA was extracted from three independent cell cultures using Qiagen RNAeasy plus kits (Qiagen 74134). *Human RNA. C*ommercially available total RNA samples were acquired from Clontech®. Each sample contained pooled RNA from 3-70 individuals. Lot numbers are indicated in parentheses: human fetal whole brain (1412624A), human adult whole brain (1402002). *Mouse RNA.* Total RNA was extracted from three to five brains of male C57BL6/J mice per age group. Brain tissue was flash frozen in liquid nitrogen, and then transferred to dry ice. Individual whole brains were pulverized with a mortar and pestle in the presence of liquid nitrogen. 30 mg of pulverized tissue was homogenized in RLT lysis buffer (Qiagen, 74134). RNA was extracted using Qiagen RNAeasy plus kit (Qiagen, 74134) according to manufacturer protocols and eluted in diethyl pyrocarbonate-treated water. *Total RNA from sorted cells*. Total RNA was extracted using TRIzol LS and isopropanol precipitation with the addition of 30 μg of GlycoBlue^®^ Coprecipitant (ThermoFisher Scientific, AM9516).

### 2.5 RT-PCR

1 μg of total RNA was primed with oligo-dT and reverse transcribed using Superscript IV according to manufacturer protocols (Invitrogen, 18091050). 2 μL of cDNA were used in a 50 μL PCR reaction containing Taq polymerase (Thermofisher, 4398881) and species-specific primers (Table 2). Cycle number was individually determined for each primer pair to amplify within the dynamic range of the PCR (Table 2). All PCRs were performed using the following conditions: 1 cycle of 95°C for 10 min, number of cycles according to Table 1 at 95°C for 30 s, 56°C for 30 s, and 72°C for 30 s, and a final cycle of 72°C for 7 min.

**Table 2.**
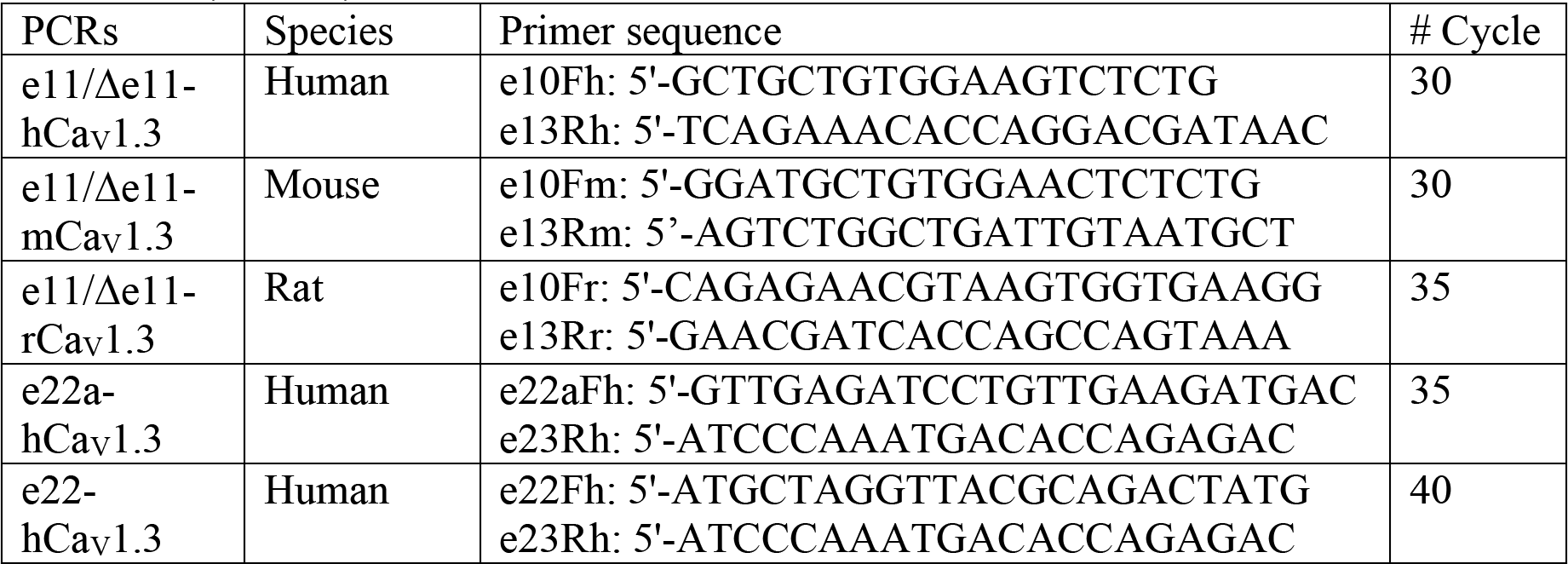
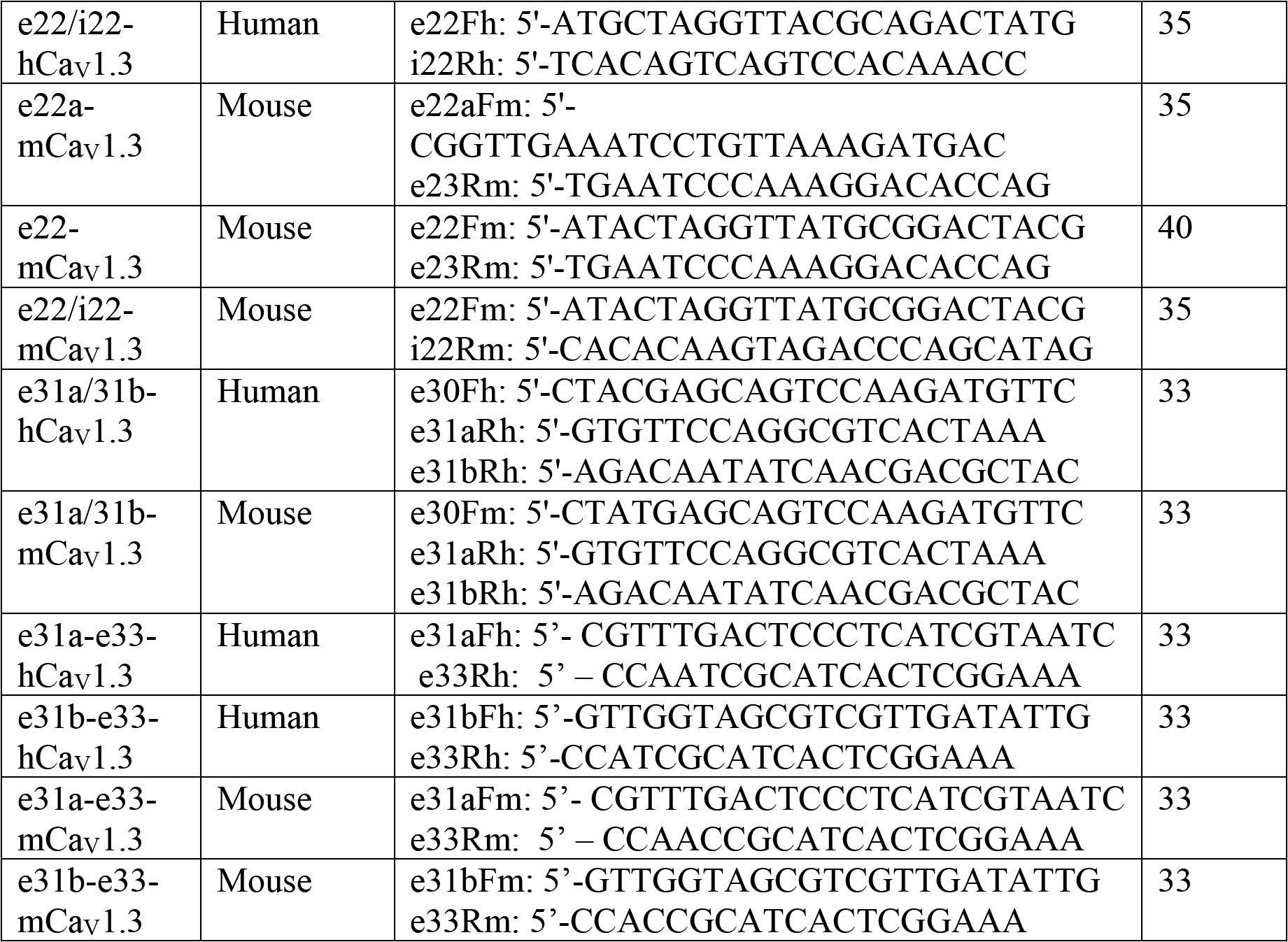
Primer sequences and cycle number for RT-PCRs performed among samples from human, mouse, and PC12 cells.

PCR products were separated in 2-3 % agarose gels and stained with ethidium bromide. Gels were imaged in unsaturated conditions using GelDoc™ XR+ Gel Documentation System (BioRad, 1708195). Densitometry analysis for each band was performed using ImageJ (Schneider et al., 2012). To control for accuracy of isoform quantification, we used reciprocal concentrations of cDNA derived from clones +e11-*Cacna1d* (Addgene, 49332) and Δe11-*Cacna1d* as reported before (Allen et al., 2017), as well as e31a- and e31b-*Cacna1d* cDNAs. The identity of each amplicon was confirmed via band extraction and sequencing. Bands were extracted using QIAquick gel extraction kit (Qiagen, 28704) according to manufacturer protocols. Extracted DNA was then either cloned using NEB PCR cloning kit (New England Biolabs, E1202S) or reamplified and purified using Zymo DNA Clean and Concentrator (Zymo research, D4064). Plasmids or PCR product were then sent to Genewiz, Inc for Sanger sequencing.

### 2.6 Electrophysiological analysis of *Cacna1d* splice variants in tsA201 cells

Δe11- and +e11-*Cacna1d* splice variants were transiently expressed in human tsA201 cells with Ca_V_β_3_ (Addgene, 26574), Ca_V_α_2_δ-1 (Addgene, 26575), and enhanced green fluorescent protein cDNAs (eGFP; BD Bioscience) as described previously (Raingo et al., 2007). Calcium channel currents were recorded 72 hours after transfection. Recordings from cells expressing either splice variant were interspersed, and the experimenter was blind to clone identity over the recording and analysis period. Clone identity was revealed post analysis. Standard whole-cell patch clamp recording was used to compare biophysical properties of both Ca_V_1.3 splice variants (Raingo et al., 2007). External solution contained: 10 mM BaCl_2_, 10 mM HEPES, 135 mM tetraethylammonium (TEA) chloride, pH adjusted to 7.2 with TEA-OH (310 mOsm). Internal solution contained: 126 mM CsCl, 10 mM EGTA, 1 mM EDTA, 10 mM HEPES, 4 mM MgATP, pH 7.2 with CsOH (290 mOsm). Barium currents were evoked by voltage-steps and currents leak subtracted on-line using a P/-4 protocol. Data were sampled at 20 kHz and filtered at 2 kHz (−3 dB). All recordings were obtained at room temperature.

### 2.7 Fluorescence activated cell sorting (FACS)

Adult *CaMKIIα;tdT* and *Gad2;tdT* mice were deeply anesthetized with isofluorane. After decapitation, brains were quickly removed and placed on a petri dish with Earl’s Balanced Salt Solution (EBSS) (Sigma, E3024) containing 21 U/mL of papain. Cerebral cortex was rapidly dissected (< 45 s), and tissue was dissociated using a modified version of Worthington Papain Dissociating System^®^ (Worthington Biochemical Corporation, LK003150). Tissue was incubated with papain for 45 min at 37° C on a rocking platform, then triturated with three sequential diameter fire-polished glass pipettes. Next, cell suspensions were centrifuged at 300 g for 5 min. After discarding supernatants, pellets were resuspended in 3 mL of EBSS containing 0.1 % of ovomucoid protease inhibitor and 0.1 % bovine serum albumin (Worthington, LK003182) to quench papain. Cell suspension was centrifuged at 270 g for 6 min and resuspended in EBSS (3 mL). tdT-expressing cells were sorted in a Sony SH800 flow cytometer using a 561-nm laser. Events were collected directly into TRIzol™ LS Reagent (ThermoFisher Scientific, 10296028). Collection was performed keeping 1:3 (v/v) sorted cell suspension: TRIzol™ LS ratio.

### 2.8 RT-qPCR

300 ng of total RNA extracted from sorted cells was primed with oligo-dT and reverse transcribed with Superscript IV First-Strand Synthesis System (ThermoFisher Scientific, 18091050) according to manufacturer instructions. RT-qPCR reactions were run on an ABI 7500 Fast Real-Time PCR system (Applied Biosystems) with the following conditions: 1 cycle 95° C for 2 min, 45 cycles (95° C for 15 s and 60° C for 1 min). Each sample from at least three different male mice per genotype (biological replicates) was run in triplicate (technical replicates). Ct values were determined by 7500 Software v2.3 (Applied Biosystems). To validate the identity of sorted cells *Gad2* and *CaMKIIα* mRNAs were quantified using TaqMan^®^ real-time PCR assays (ThermoFisher Scientific) with the following probes Mm00484623_m1 and Mm00437967_m1. mRNA levels were normalized to *Gapdh* using the probe Mm99999915_g1. Relative quantification of gene expression was performed with the ΔΔ-Ct method (Livak and Schmittgen, 2001).

### 2.9 Statistical analysis

Data are presented as mean ± one standard error of the mean (s.e.m.). Student’s t-tests and one-way analysis of variance (ANOVA) were performed with SPSS (IBM). Bonferroni correction was used to adjust for multiple comparisons. We used G*power (http://www.gpower.hhu.de/) to perform power analyses. To detect differences in mean values between two independent groups with 80 % power and a confidence interval of 95 %, we determined a sample-size of 3-5 was needed for each group.

## 3. RESULTS

### 3.1 Inclusion of e11 in the final *Cacna1d* mRNA is upregulated during postnatal brain development

The DI-II linker is highly variable across the major mammalian Ca_V_α_1_ subunits (Catterall, 2011; Lipscombe et al., 2013). This cytoplasmic region is important for membrane trafficking, interaction with Ca_V_β subunits and other regulatory proteins, coupling to signaling cascades, and fine-tuning of biophysical properties of voltage-gated calcium channels (Catterall, 2000). *Cacna1d* pre-mRNA contains a cassette exon or e11 (*GRCm38/mm10*, chr14: 30,122,938-30,136,237) that originates two Ca_V_1.3 splice variants with differences in the DI-II linker (Δe11 and +e11-*Cacna1d*) (Zuccotti et al., 2011). Exon 11 encodes for a 21-amino acid sequence located 47 amino acids downstream of the alpha-interaction domain (AID) in Ca_V_1.3 (Fig. 1A). Exon 11 is conserved among mammalian orthologs (see Vertebrate Multiz Alignment & Conservation, 100 Species, track of the UCSC Genome Browser. Fig. 1B). To determine the temporal pattern of e11 during postnatal brain development, we quantified the relative amounts of Δe11- and +e11-*Cacna1d* in mouse whole brains from multiple ages. We found that at postnatal day 0 (P0), inclusion of e11 is rare and drastically increases between P9-P18, but stabilizes after P30 through adulthood (Fig. 1C). To calibrate the relative amounts for Δe11- and +e11-*Cacna1d*, we performed non-saturating PCR reactions on samples containing pre-determined reciprocal amounts of plasmid cDNA from *Cacna1d* clones with and without e11. Note that predicted and experimental values are similar (Fig. 1D). Next, we performed RT-PCR for Δe11- and +e11-*Cacna1d* in RNA samples from fetal and adult human whole brain (fWB and aWB respectively, Fig 1E). We found that e11-*Cacna1d* is also rare in fWB, whereas it dominates in aWB (Fig. 1E and (Charizanis et al., 2012)). Our data show that inclusion of e11 in *Cacna1d* is upregulated during postnatal brain development in mouse and humans.

### 3.2 Excitatory pyramidal neurons preferentially express +e11-*Cacna1d*, whereas inhibitory interneurons express mostly Δ11-*Cacna1d* splice variants

We next aimed to determine what neuronal types express Δe11- and +e11-*Cacna1d* splice variants in adult brain. Previous studies have shown that INs and PNs express opposite patterns of alternative splicing for several genes (Tasic et al., 2016; Zhang et al., 2016). Therefore, we tested if Δe11 and +e11-*Cacna1d* splice variants show differential expression in INs and PNs of adult mouse cortex. In order to do this, we performed FACS in genetically labeled neuronal populations coupled to RT-PCR (Fig. 2A). INs and PNs were labeled using the *Cre-loxP* system (see methods). INs were labeled with tdT and *Gad2* promoter, whereas PNs with tdT and *CaMKIIα* promoter. Cortical tissue from *Gad2;tdT* and CaMKIIα;tdT mice was dissociated, and RNA was extracted from tdT expressing cells from both tissues (Gad2^+^ and CaMKIIα^+^, respectively). We next validated the extracted RNA by quantifying *Gad2* and *CaMKIIα* mRNA in both samples using TaqMan gene expression assays. As predicted, RNA from Gad2^+^ cells contained significantly larger amounts of *Gad2* mRNA than RNA extracted from CaMKIIα^+^ cells (Mean 2^−ΔΔCt^ ± SE: Gad2^+^ = 29.33 ± 6.33, n = 3; CaMKIIα^+^ = 1.0 ± 0.15, n = 3. Student’s t-test, p = 0.009. Fig. 2B, *left panel*). Similarly, CaMKIIα^+^ cells expressed larger amounts of *CaMKIIα* mRNA relative to Gad2^+^ cells (Mean 2^−ΔΔCt^ ± SE: Gad2^+^ = 1.00 ± 1.10, n = 3; CaMKIIα^+^ = 76.37 ± 36.16, n = 3. Student’s t-test, p = 0.008. Fig. 2B, *right panel*). We next quantified the levels of Δe11- and +e11-*Cacna1d* splice variants in RNA derived from Gad2^+^, CaMKIIα^+^ cells, and whole brain cortex. We found that Δe11-*Cacna1d* is dominant in INs, whereas +e11-*Cacna1d* dominates in PNs, interestingly this differs from whole cortex where both splice variants are expressed at similar levels (Mean % e11 ± SE : cortex = 44.59 ± 4.3, n = 3; Gad2^+^ = 0.62 ± 0.41, n = 3; CaMKIIα^+^ = 84.5 ± 2.09, n = 3. ANOVA, F_2,8_ = 229.67, p < 0.0001. Fig. 2C). Post hoc analysis with Bonferroni correction showed that all means are statistically significant. Our results strongly suggest that INs express Δe11, whereas PNs include e11 in the final *Cacna1d* mRNA.

**Figure 2.**
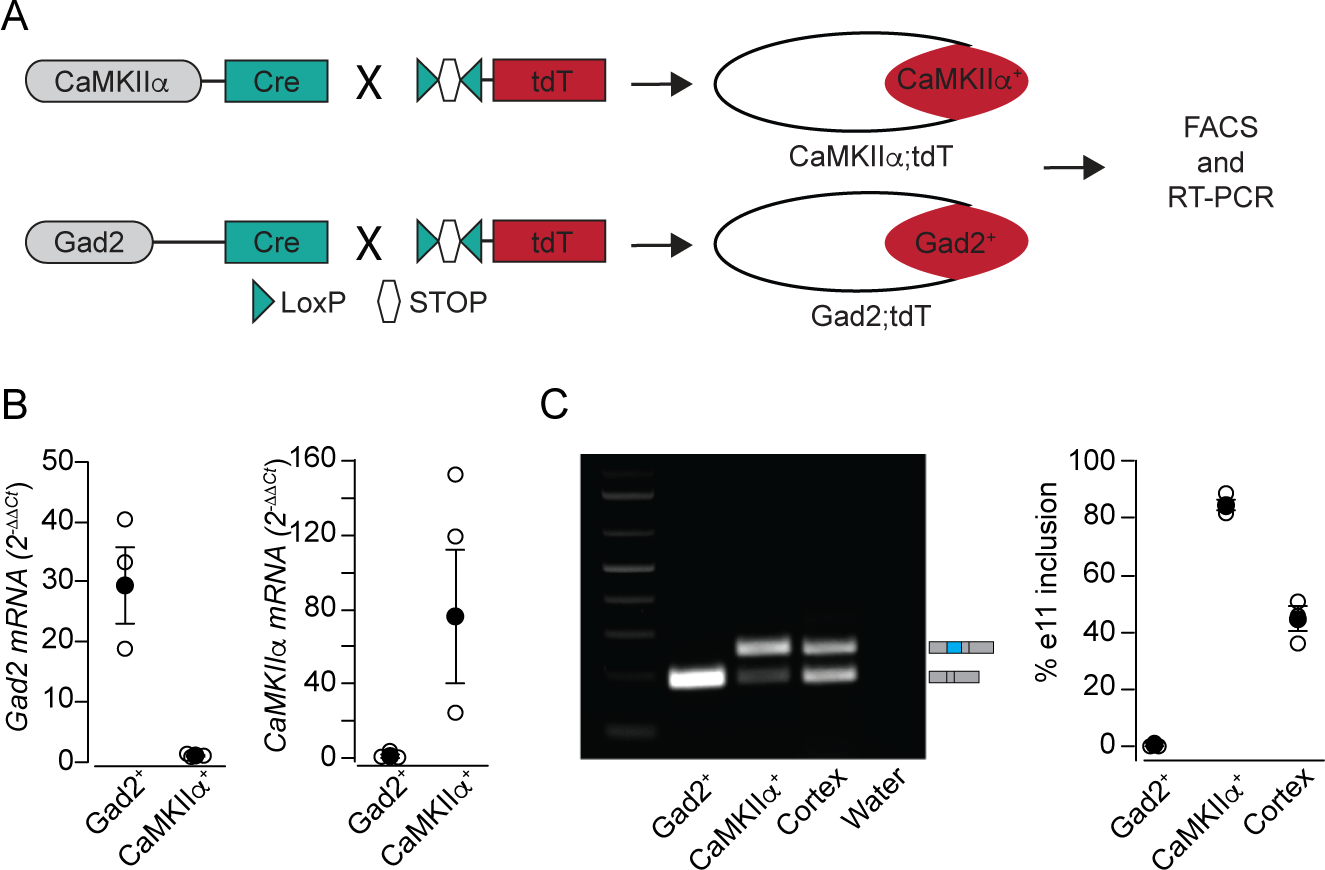
Cell-specific expression of Δe11- and +e11-*Cacna1d* splice variants. **A**) Schematic showing the workflow to extract RNA of excitatory PNs and inhibitory GABAergic INs. *Gad2* and *CaMKIIα* promoter in PNs and GABAergic INs drive the expression of *Cre*-recombinase, which in turn excises a STOP codon flanked by LoxP sites in an allele containing tdT. This results in expression of tdT in CaMKIIα^+^ and Gad2^+^ cells. These cell populations were purified using FACS, and total RNA was extracted to quantify *Cacna1d* splice variants. **B**) RNA extracted from sorted cells was validated by quantifying *Gad2* and *CaMKIIα* mRNA. **C**) *Left panel*, representative RT-PCR gel performed to quantify +e11- and Δe11-*Cacna1d* splice variants in sorted cells and whole cerebral cortex. *Right panel*, quantification of e11 alternative splicing in Gad2^+^ and CaMKIIα^+^ cells, as well as cerebral cortex. All data are shown as mean ± s.e.m (solid symbols), and individual biological replicates (empty symbols).

### 3.3 Differentiation by nerve-growth factor upregulates +e11-*Cacna1d* in PC12 cells

During brain development, embryonic neural progenitor cells differentiate into a wide diversity of neurons to form mature neuronal circuits (Huang and Reichardt, 2001). Neuronal differentiation is accompanied by the determination of splicing programs, which underlie cell-type specific functions (Baralle and Giudice, 2017). Progenitor cells are exposed to factors that induce differentiation into defined cell-types during brain development. NGF is a classical growth factor important for survival, neurite outgrowth, and sprouting of sensory, sympathetic and central cholinergic neurons (Sofroniew and Howe…, 2001). Upon exposure to NGF, PC12 cells develop a neuronal-like phenotype. Here we assessed if NGF influences alternative splicing of the *Cacna1d* pre-mRNA in PC12 cells. We found that undifferentiated PC12 cells (UD) expressed lower levels of +e11-*Cacna1d* relative to PC12 cells differentiated with NGF (DF) (Mean ± SE % e11. UD = 24.5 ± 0.5, n = 3; DF = 46.6 ± 1.5, n = 3. Student’s t-test, p = 0.0001, Fig. 2A). These results show that NGF induces inclusion of e11. This event nicely correlates with inclusion of e11 in adult brain and PNs, suggesting the presence of shared mechanisms to induce inclusion of e11 in the final *Cacna1d* mRNA.

### 3.4 Comparison of biophysical properties between Δe11 and +e11-*Cacna1d* splice variants in mammalian expression systems

Previous studies comparing currents of Δe11-Ca_V_1.3 and +e11-Ca_V_1.3 channels expressed in oocytes together with auxiliary subunits, Ca_V_β_1_ and Ca_V_α_2_δ-1, showed no differences in gating between these two splice variants. Here we compared several biophysical properties between Δe11- and +e11-*Cacna1d* splice variants using the mammalian expression system, tsA201 cells and in the presence of the most abundant Ca_V_β subunit in the brain, Ca_V_β_3_. Using whole-cell voltage-clamp, we measured current density (Fig. 4A **and 4B**), voltage-dependent activation and inactivation (Fig. 4C **and 4D**), as well as open state inactivation (Fig. 4E). We used barium as charge carrier to prevent Ca^2+^-dependent inactivation. No statistically significant differences were found in any of the parameters that describe these biophysical properties (see table 3), suggesting that both Δe11- and +e11-*Cacna1d* splice variants produce fully functional Ca_V_1.3 channels with similar biophysical properties. Our findings also confirm previous studies in oocytes.

**Table 3.**
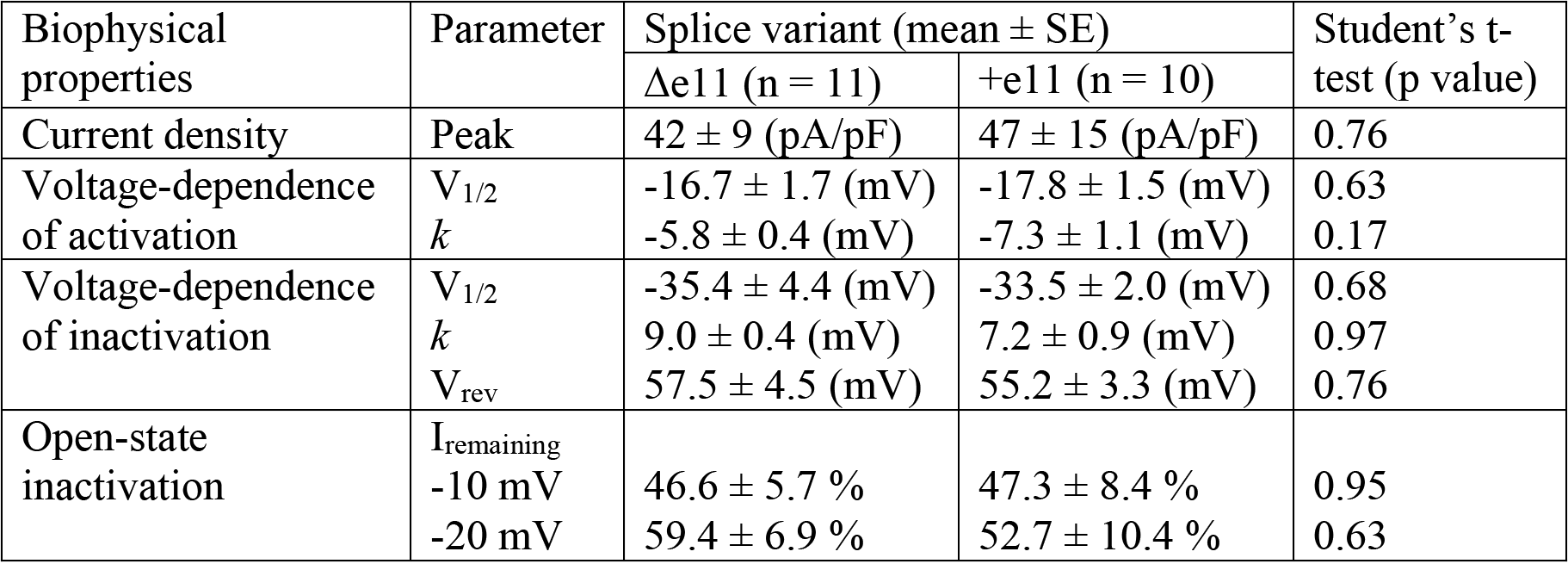
Biophysical properties of +e11-*Cacna1d* and Δe11-*Cacna1d* splice variants expressed in tsA201 cells.

**Figure 3.**
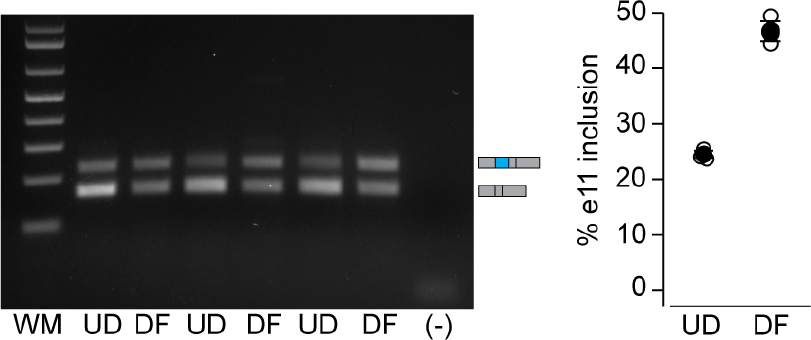
Differentiation of PC12 cells into a neuron-like phenotype induces inclusion of e11 in the *Cacna1d* mRNA. *Left panel*: Representative agarose gel image of RT-PCR from undifferentiated (UD) and NGF-differentiated (DF) PC12 cells. Rat specific primers were used in RT-PCR performed from three independent PC12 cultures. *Right panel:* Summary data from UD and DF PC12 cells. Data are mean (black circles) ± s.e.m. % e11. UD = 24.5 ± 0.5, DF = 46.6 ± 1.5. Empty circles represent individual values. Empty circles represent individual values, and solid circles represent means.

**Figure 4.**
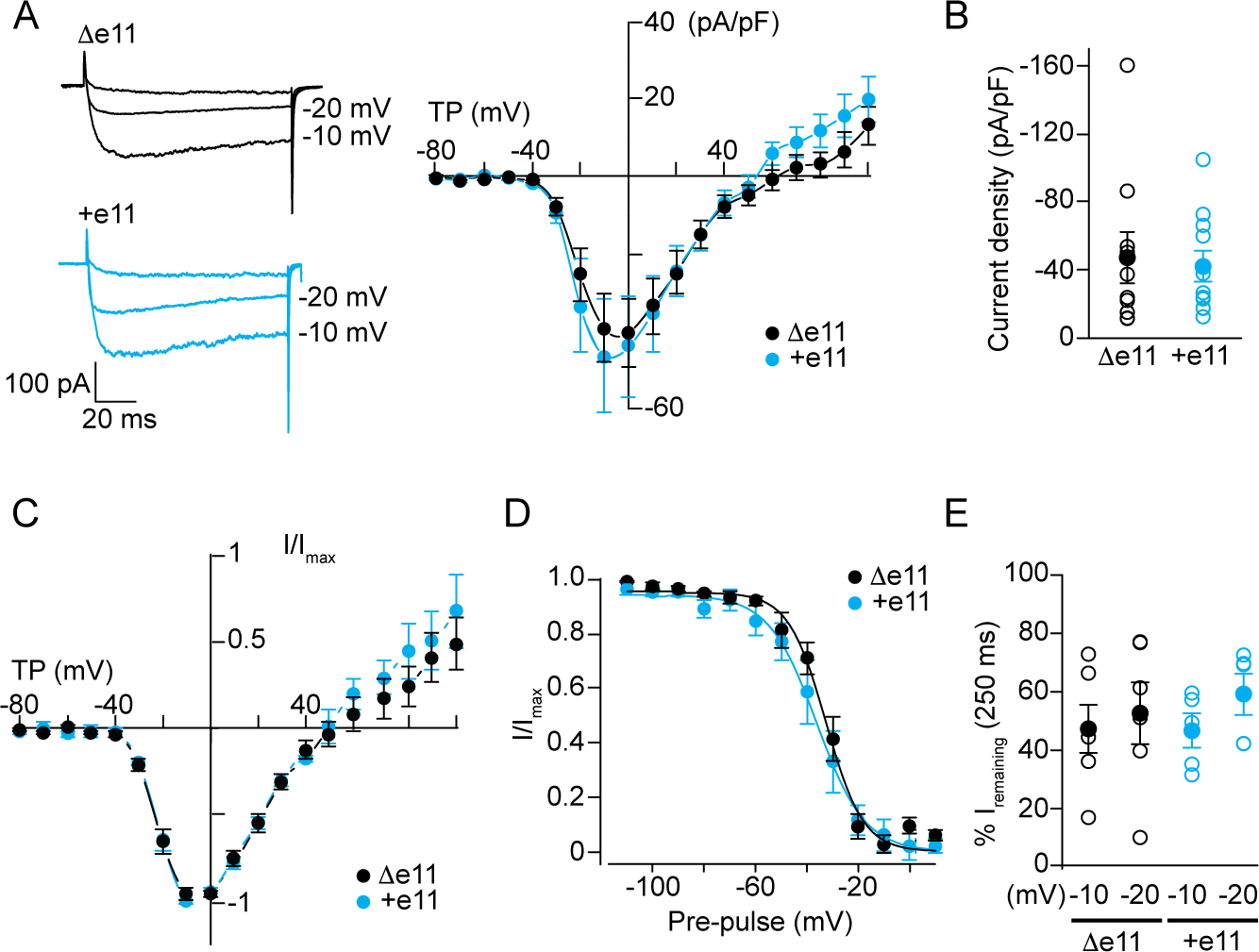
Comparison of biophysical properties between +e11- and Δe11-*Cacna1d* splice variants. **A**) *Left panel*, representative traces of whole-cell currents recorded 72 hrs after transfection of tsA201 cells with clones expressing each splice variant together with Ca_V_β_3_ and Ca_V_α_2_δ-1. Barium was used as the charge carrier. Currents were evoked by voltage steps of 10 mV increments applied from a holding potential of −80 mV. Currents recorded from cells expressing Δe11- and +e11-*Cacna1d* cDNAs are shown. *Right panel*, current-voltage (I-V) relationships of Δe11- and +e11-*Cacna1d* splice variants are compared. Peak current densities are plotted against test potential (TP). Currents were elicited by step depolarizations from −80 mV to 80 mV in 10 mV increments from a holding potential of −80 mV. I-V relationships were fit to data from individual cells using the product of the Boltzmann function and a straight line for estimates of activation mid-point (V_1/2_), slope factor (k) and reversal potential (V_rev_) (See table 3). **B**) Peak current densities evoked at −10 mV from individual cells (open symbols) expressing Δe11- and +e11-*Cacna1d* splice variants are shown together with averaged current density and mean ± s.e.m. (solid symbols). **C**) Normalized I-V from the same cells as in **A**. **D**) Steady state inactivation relationships from normalized peak current amplitudes at −10 mV following 3 s pre-pulse steps applied in 10 mV increments to potentials between −110 mV and 10 mV. Boltzmann functions were fit to each data set and mean ± SE values for both splice variants are shown. **E**) Open-state inactivation was measured as percentage of current that remains after 250 ms of depolarizing pulse. Remaining current was measured at −10 and −20 mV from a holding potential of −80 mV. Data are presented as mean ± s.e.m.

### 3.5 Inclusion of exon 22 in *Cacna1d* pre-mRNA is upregulated in human brain development

The *Cacna1d* pre-mRNA contains a tandem of mutually exclusive exons that encode for S2 sequences of DIII in Ca_V_1.3, e22a and e22 (Fig. 5A). Genome tracks show high peaks of conservation corresponding to e22a (*GRCm38/mm10*, chr14: 30,107,786-30,107,845. Fig. 5B) and e22 (*GRCm38/mm10*, chr14: 30,107,653-30,107,712. Fig. 5B). Previous reports have shown that e22a, but not e22, is exclusively used in adult rat brain, whereas e22 is used in auditory hair cells of rat, trout and chicken (Kollmar et al., 1997; Ramakrishnan et al., 2002). However, alternative splicing of e22a and e22 has not been assessed in human or mouse brain. To do this, we performed RT-PCR in fWB and aWB human tissues, as well as brain tissue from newborns and adult mice (P0 and P180, respectively). We used primers that anneal to sequences within e22a and e22 (Fig. 5A). We found that e22a is used in P0 and P180 mouse brain, as well as fetal and adult human brain (Fig. 5B, lanes 2, 4, 8, and 10). We also found that e22 is not present in fetal human brain, but it is slightly used in adult human brain (Fig. 5C, lanes 9 and 11). Surprisingly, we were unable to identify e22 in either P0 or P180 mouse brains (Fig. 5C, lanes 3 and 5). To validate that our primers were performing properly, we PCR-amplified from genomic DNA of mouse and human samples using the same forward primer annealing to e22 (e22Fm) and a reverse primer annealing to an intronic region downstream of e22 (i22Rm). We observed robust amplification using this set of primers, suggesting that e22Fm primer is capable of annealing the predicted cDNA target (Fig. 5A **and** Fig. 5C, lanes 6 and 12). This data show that e22 is present in aWB, but not in fWB. In contrast, e22 is not present in either P0 or P180 mouse tissue.

**Figure 5.**
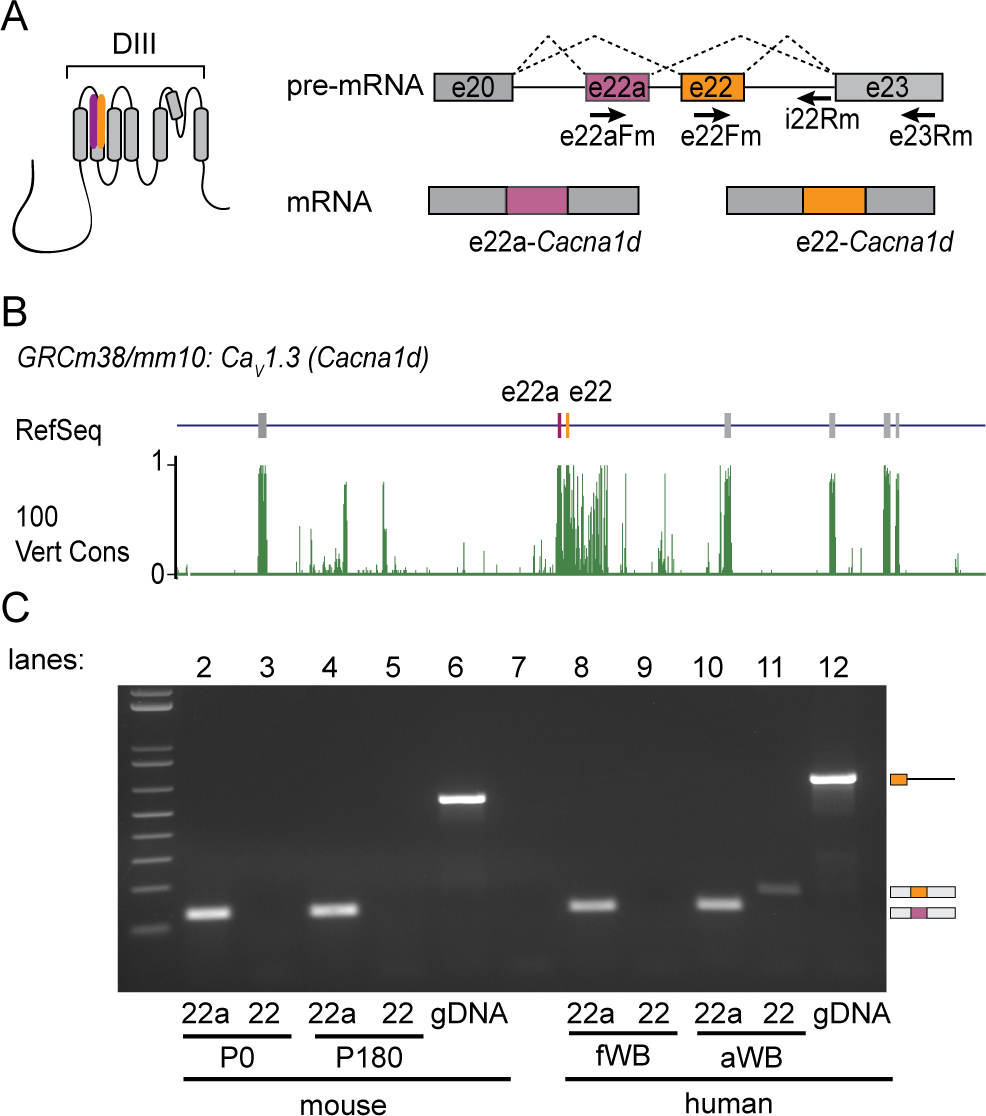
Comparison of alternative splicing of e22a and e22 in *Cacna1d* over development of mouse and human brain. **A**) *Left panel:* DIII of Ca_V_α_1_ pore-forming subunit. Highlighted in maroon and yellow is the putative location of sequences encoded by e22a and e22 in DIII respectively. *Right panel:* Schematic representation of alternative splicing of the mutually exclusive exons 22a and 22. Exons are shown as boxes and intronic regions as black lines. Putative location of PCR primers used to amplify human and mouse splice variants and genomic DNA (gDNA) are shown as black arrows. **B**) Visualization of RefSeq Genes and 100 vertebrates Basewise Conservation by PhastCons tracks using UCSC Genome Browser. The region of the mouse *Cacna1d* gene (GRCm38/mm10, chr14: 30,101,395-30,112,778). This region shows exons 21, 22a, 22, 23, 24, 25, and 26. In the RefSeq track, exons are depicted with vertical bars and introns with horizontal lines. Maroon and yellow vertical bars denote locations of e22a and e22 respectively. The conservation track displays the PhastCon scores from 0 to 1. Both exons are highly conserved across vertebrate species. **C**) RT-PCR products for e22a and e22 in human and mouse whole brain across development. E22a is present in both P0 (lane 2) and P180 (lane 4) mouse brains. E22 is not present in either P0 (lane 3) or P180 (lane 5) mice. E22a is present in fWB (lane 8) and aWB (lane 10). Exon 22 was not detected in fWB (lane 9), but is present in aWB (lane 11). gDNA from mouse (lane 6) and human (lane 12) showed robust amplification using primers to detect e22 in mouse and human brain (see Table 1).

### 3.4 Alternative processing of exons 31a, 31b and 32 of*Cacna1d* pre-mRNA during postnatal brain development

Exons 31a (*GRCm38/mm10*, chr14: 30,107,786-30,107,845) and 31b (*GRCm38/mm10*, chr14: 30,089,296-30,089,380) are a pair of mutually exclusive exons in the *Cacna1d* pre-mRNA that generate splice variants with different sequences in S3 DIV (Fig. 6A). However, the pattern of 31a and 31b exon usage during brain development has not been fully determined. Here we tested if e31a and e31b inclusion in the final *Cacna1d* mRNA changes throughout development of human and mouse brain tissues. To assess this, we PCR-amplified using a common forward primer located in e30 and two reverse primers located in either e31a or e31b (Fig. 6A). We observed that inclusion of e31a is lower in P0 mouse brains than in P180 mouse brains (Data are mean ± SE % e31a inclusion: P0 = 3.8 ± 1.2, n =3; P180 = 53.8 ± 1.2. Student’s t-test, p < 0.0001. Fig. 6B). Similar results were found in human brain, e31a is rarely used in fetal brain, but increases in adult whole brain (Fig. 6C, *left panel*). To validate that the set of primers and PCR conditions amplify e31a- and e31b-containing *Cacna1d* splice variants with similar efficacy, we performed PCR using reciprocal amounts of plasmid DNA containing either e31a or e31b. Our experimental values were similar to the predicted values (Fig. 6C, *right panel*). Our results suggest that inclusion of 31a increases during brain development in both mouse and human brain. Exons 31a and 31b can also be spliced in together, e31a:31b (Fig. 6A). Inclusion of e31a:e31b is predicted to produce a shift in the reading frame, introducing a premature stop codon. We found that this splice variant is only detectable prior to P9 and in fWB. Our results show that the e31a:e31b splice variant is repressed at early stages of brain development in both mouse and human brain.

**Figure 6.**
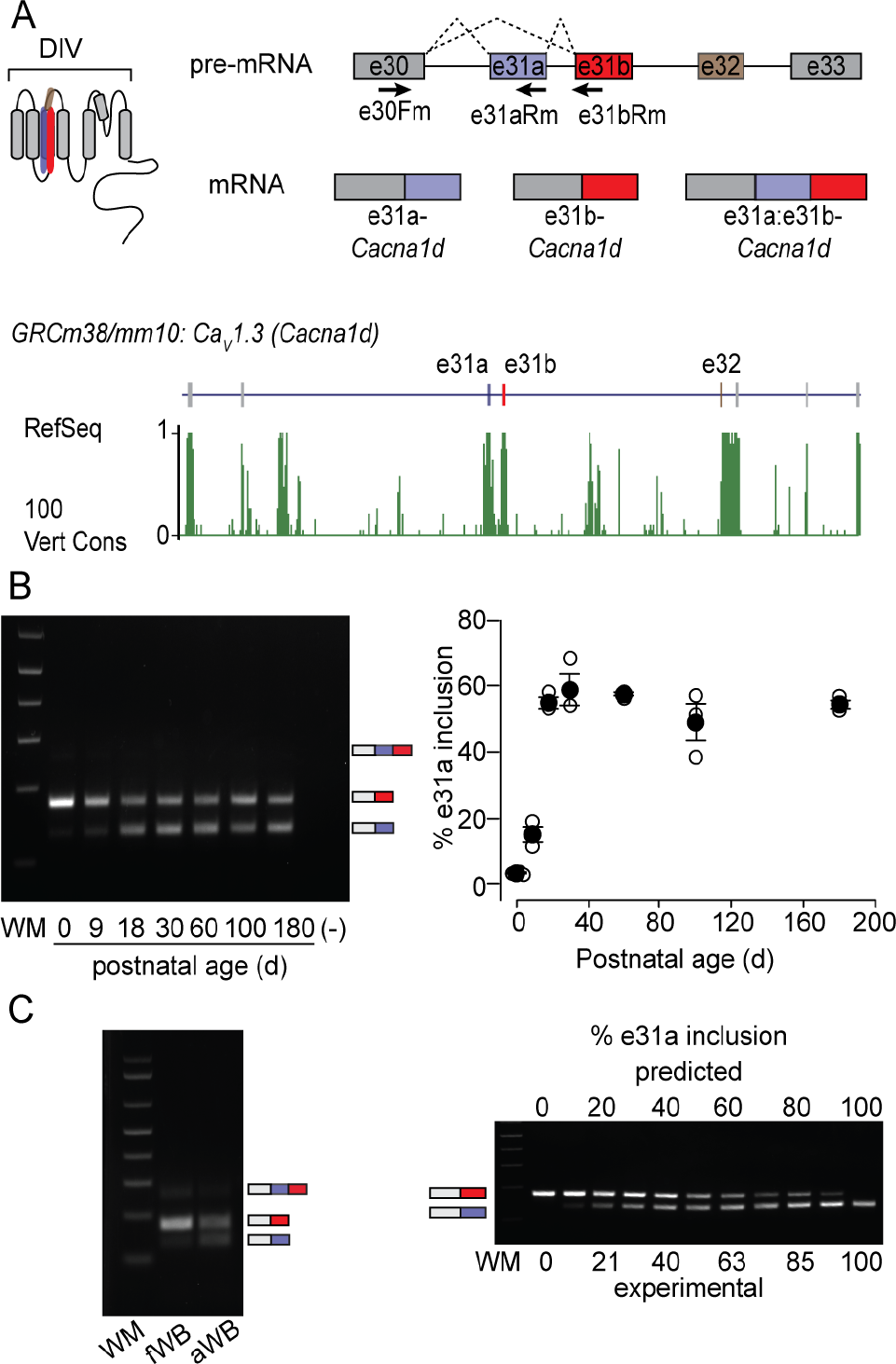
Alternative splicing of e31a and e31b of the *Cacna1d* pre-mRNA during brain development. **A**) *Upper panel:* DIV of Ca_V_α_1_ pore-forming subunit. Highlighted in purple and red are the putative locations of exon 31a and 31b, respectively. Highlighted in brown is the putative location of the cassette exon 32. Amplification of the region from e30 to either e31a or e31b results in three splice variants containing e31a, e31b or a fusion of both exons (e31a:e31b). Boxes represent exons, horizontal lines represent intronic regions, and black arrows represent the approximate location of PCR primers used to amplify human and mouse cDNAs from whole brain. *Lower panel:* Visualization of RefSeq Genes and 100 vertebrates Basewise Conservation by 810 PhastCons tracks using UCSC Genome Browser for the mouse *Cacna1d* gene (*GRCm38/mm10*, chr14:30,077,665-30,097,455). The region shown contains exons 29, 30, 31a, 31b, 32, 33, 34, and 35. In the RefSeq track, exons are denoted by vertical bars and introns by horizontal lines. Predicted locations of e31a, e31b, and e32 are shown with purple, red and brown vertical bars, respectively. The conservation track displays the PhastCon scores from 0 to 1. All three exons are highly conserved across vertebrates. **B**) Representative gel of RT-PCR to detect e31a- and e31b-*Cacna1d* splice variants in mouse brain at different postnatal ages. Mouse-specific primers were used. *Right panel:* RT-PCRs from mouse brains (3 per age group). Data are mean (black circles) ± SE % of e31a. P0 = 3.3 ± 0.3, P9 = 15.1 ± 2.2, P18 = 54.9 ± 1.6, P30 = 58.8 ± 4.8, P60 = 57.5 ± 0.62, P100 = 49.0 ± 5.5, P180 = 54.4 ± 1.2. Empty circles indicate individual values. (−) denotes negative PCR control where cDNA was replaced with water. **C**) *Left panel:* Representative gel of RT-PCR from fWB and aWB to detect e31a- and e31b-*Cacna1d* splice variants. *Right panel:* Calibration curve with reciprocal amounts of e31a and e31b cDNA template shows similar PCR efficiency of e31aR and e31bR primers. Predicted values indicate the amount of e31a template relative to 31b template added to PCR master mix.

Alternative splicing of the cassette exon e32 (*GRCm38/mm10*, chr14: 30,082,952-30,082,997, Fig. 6A **and** 7A) produces *Cacna1d* splice variants with modifications in the extracellular loop between S3 and S4 of DIV (Fig. 7A). We analyzed the developmental expression of e32 in samples from human and mouse brain. Because e32 can be spliced with either e31a or e31b, we performed two independent PCRs, one amplifying from e31a through e33 (Fig. 7A), and a second one amplifying from e31b through e33 (Fig. 7C). Amplification from e31a through e33 resulted in four detectable splice variants, e31a:e31b:e32, e31a:e31b:Δe32, e31a:e32, and e31a:Δe32 (Fig. 7A **and** Fig. 7B) in both mouse and human brain. Analysis of mouse brain tissue showed that the four 31a-containing *Cacna1d* splice variants mentioned above were present at P0, but the e31a:e32 *Cacna1d* splice variant dominates in adult whole brain (Fig. 7B). We observed something similar in human brain, e31a-containing *Cacna1d* splice variants that also include e31b and e32 (e31a:31b:e32) are more abundant in fWB than in aWB, whereas e31a-containing *Cacna1d* splice variants that only include e32 (e31a:e31b:e32 and e31a:e32) are more abundant in adult brain than in fWB (Fig. 7B, right). PCR-amplification from e31b through e33 allowed the detection of two splice isoforms, e31b:e32 and e31b:Δe32 (Fig. 7D). In mouse brain we found that e32 is slightly repressed over brain development (Fig. 7D, *left panel*). In human tissue, we e31b:e32 is expressed at higher levels in fWB relative to aWB (Fig. 7D, *right panel*). Our results show that: *i*) inclusion of e32 is upregulated in adult brain when e31a is included, but it is slightly repressed when e31b is used, and *ii*) splice variants containing e31a:e31b are downregulated in adult brain, confirming the results above.

**Figure 7.**
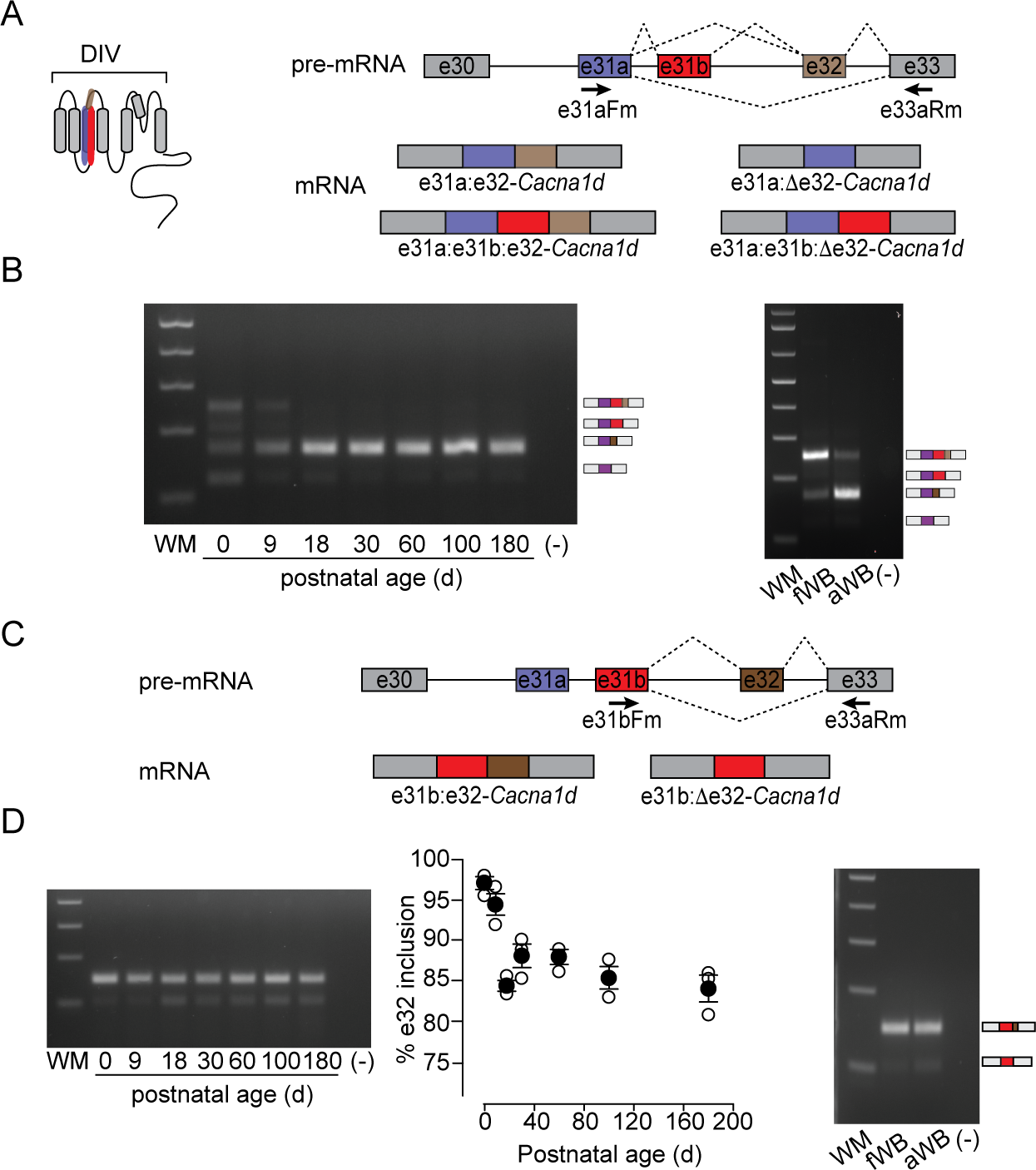
Alternative splicing of e32 in the *Cacna1d* pre-mRNA throughout brain development. **A**) *Left panel:* Schematic of DIV of Ca_V_α_1_ pore-forming subunit. *Right panel:* Amplification of from e31a to e33 can yield four different splice variants: e31a:e32-*Cacna1d*, e31a:Δ32-*Cacna1d*, e31a:e31b:e32-*Cacna1d*, and e31a:e31b:Δ32-*Cacna1d*. Arrows indicate the estimated location of primers. **B**) *Left panel*, RT-PCR from mouse brains at different postnatal ages. The relative amounts of each of the four splice variants are (mean ± SE %). e31a:e32: P0 = 22.6 ± 7.1, P9 = 83.4 ± 3.3, P18 = 95.7 ± 0.8, P30 = 98.8 ± 0.1, P60 = 97.4 ± 0.3, P100 = 97.6 ± 0.2, P180 = 98.0 ± 0.3. e31a:Δ32: P0 = 33.3 ± 6.7, P9 = 7.4 ± 0.7, P18 = 4.2 ± 0.8, P30 = 1.2 ± 0.1, P60 = 2.6 ± 0.5, P100 = 2.3 ± 0.2, P180 = 2.0 ± 0.3. e31a:e31b:e32: P0 = 39.6 ± 4.5, P9 = 9.2 ± 2.8. e31a:e31b:Δ32: P0 = 4.5 ± 1.6. *Right panel*, Representative gel of RT-PCR from fWB and aWB from human. **C**) Splicing pattern for e32 when e31b in the presence of e31b. Arrows indicate the estimated location of the primers used for PCR amplification. **D**) *Left panel*, RT-PCRs from mouse brains (3 per age group). Representative gel and mean (solid symbols circles) ± SE of % e32 inclusion in whole mouse brain at different postnatal ages. P0 = 97.2 ± 0.8, P9 = 94.5 ± 1.4, P18 = 84.4 ± 0.7, P30 = 88.1 ± 1.1, P60 = 88.0 ± 0.9, P100 = 85.3 ± 1.4, P180 = 84.0 ± 1.6. Empty circles indicate individual values. *Right panel*, Representative RT-PCR gel of fWB and aWB in human.

## 4. DISCUSSION

The *Cacna1d* pre-mRNA contains exons that alter sequences in DI, I-II linker, DIII and DIV and C-terminus. In this study we focused on alternative spliced exons that generate splice variants with structural differences in the I-II linker, DIII and DIV of Ca_V_1.3. These exons are cassette e11, mutually exclusive exons 22a/22, mutually exclusive 31a/31b, and the cassette exon 32. The function of these exons is not fully understood. However, determining their temporal and cell-specific expression is a step towards understanding their role in Ca_V_1.3 channel activity.

### 4.1 Regulation of *Cacna1d* pre-mRNA alternative splicing during postnatal brain development

Identification and characterization of splicing factors have helped explain exon usage during development. Previous studies have suggested that three different splicing factors can influence alternative splicing of e11 in the *Cacna1d* pre-mRNA. Knock-out mice for the splicing factors muscleblind-like splicing regulator 1 and 2 (Mbnl1 and Mbnl2) and RNA binding protein fox homologue 1 (*Rbfox1*) show decreased inclusion of e11 in whole brain (Gehman et al., 2011; Charizanis et al., 2012). Interestingly, expression of these splicing factors increases over development, and have been shown to regulate the transition from fetal to adult splicing patterns. Mbnl1 and Mbnl2 specifically repress embryonic stem cell splicing patterns and promote differentiated neuronal cell splicing patterns (Lin et al., 2006; Kalsotra et al., 2008; Venables et al., 2013), while *Rbfox1* has been shown to regulate both splicing and transcriptional networks in human neuronal development (Fogel et al., 2012; Weyn-Vanhentenryck et al., 2014; Hamada et al., 2015). Developmental changes in L-type calcium channel function have also been observed (Ghosh et al., 1994; Flavell and Greenberg, 2008). It is possible that splicing factors influence calcium dynamics during development by inducing the proper alternative splicing of voltage-gated calcium channel pre-mRNAs.

Our results show that e22a dominates in both fetal and adult whole brain of mice and human. Exon 22 was not identified in fetal human brain, but was present at low levels in adult human brain, suggesting that inclusion of e22 increases over development. The splicing factors that regulate e22a/e22 in *Cacna1d* pre-mRNA have not been fully characterized. Alternative splicing of the locus e31a/e31b/e32 originates up to six distinct *Cacna1d* splice variants. In cases where e31a:e31b is included, a change in the open reading frame is likely to occur, which is predicted to produce a premature stop codon. We also found that skipping of e32 occurs in the presence e31b, whereas inclusion of e32 occurs in the presence of e31a. The regulation of this complex alternative splicing event is not fully understood; however, analysis of RNA isolated by crosslinking immunoprecipitation (HITS-CLIP) shows unique binding tags for the splicing factors NOVA1, NOVA2, Rbfox1, Rbfox2 and Rbfox3 (Ule et al., 2003; Weyn-Vanhentenryck et al., 2014). This raises the possibility that these splicing factors co-regulate splicing of e31a/e31b and e32 in the *Cacna1d* pre-mRNA.

### 4.2 Functional implications and cell-specific expression Δe11- and +e11-*Cacna1d* splice variants

Our studies show that Δe11 and +e11-*Cacna1d* splice variants generate fully functional Ca_V_1.3 channels with similar biophysical properties in mammalian expression systems. Although these results fail to point at functional differences between the two splice variants, the prime localization of e11 in the I-II linker suggests a role in membrane trafficking, coupling to signaling cascades or interaction with cytoplasmic proteins such as Ca_V_β subunits. Furthermore, 30 % of amino acids in e11 are positively charged, and high-throughput proteomic studies suggest the presence of a conserved arginine subject to methylation and a serine subject to phosphorylation (Hornbeck et al., 2014). Arginine methylation has been shown to regulate cell surface expression and overall activity of other ion channels (Beltran-Alvarez et al., 2013; Kim et al., 2016). Future studies considering these observations will be necessary to determine the function of e11. Finally, we have demonstrated that +e11-*Cacna1d* is primarily expressed in PNs, whereas Δe11-*Cacna1d* dominates in GABAergic INs of cortex. Our results point at a system to study the functional relevance of this exon.

### 4.3 Functional implications of splice variants within DIII and DIV of Ca_V_1.3

The *Cacna1d* gene has a similar genomic arrangement to the *Cacna1c* gene, as a result several splicing events are common to both pre-mRNAs. Therefore, functional studies on *Cacna1c* splice variants could provide some insights into the function of splice variants for *Cacna1d*. The *Cacna1c* pre-mRNA contains a mutually exclusive pair of exons similar to e22a and e22 in *Cacna1d*, e21 and e22. Splice variants containing e21 in *Cacna1c* show reduced sensitivity to DHPs relative to e22-including splice variants (Soldatov et al., 1995; Zühlke et al., 1998). Previous studies comparing brain Ca_V_1.3 channels (contains e22a) relative to heart Ca_V_1.2 channels (contains e22) nicely demonstrated that Ca_V_1.3 is less sensitive to DHP (Xu and Lipscombe, 2001). It would be interesting to see if splicing of e22a/e22 also modifies the sensitivity of Ca_V_1.3 channels to DHPs.

Interestingly, exons 31a, 31b and e32 in the *Cacna1d* pre-mRNA undergo alternative processing similar to e31, e32 and e33 in the *Cacna1c* pre-mRNA (Xu and Lipscombe, 2001). Alternative splicing of e31, e32 and e33 has been shown to also affect the sensitivity of Ca_V_1.2 channels to DHP (Soldatov et al., 1995; Zühlke et al., 1998). However, the few functional studies regarding e31a, e31b and e32 in *Cacna1d* shed no differences in biophysical properties or DHP sensitivity (Xu and Lipscombe, 2001). Suggesting that these exons have unique roles in Ca_V_1.3 channels. Understanding the functional role of these exons is a future endeavor.

### 4.4 Regulation of alternative splicing the *Cacna1d* pre-mRNA by growth factors

Alternative splicing is sensitive to developmental cues carried out by neurotrophic factors such as epidermal growth factor (Zhou et al., 2012). Furthermore, several studies have shown that signaling by growth factors regulates the activity and expression of splicing factors (Liu et al., 2003). In this work, we report that differentiation mediated by NGF induces a splice program for *Cacna1d* similar to adult whole brain and excitatory neurons. Although the exact mechanism of this is yet to be determined, our data suggest that PC12 cells contain the elements necessary to link NGF to alternative splicing. Further work in both cell lines and primary cultures is needed to obtain more mechanistic insights into how NGF controls alternative splicing of the *Cacna1d* pre-mRNA.

## CONFLICT OF INTEREST

The authors declare no conflict of interest.

## AUTHOR CONTRIBUTIONS

LB, BA, SH, AM performed research. LB, FT and AA designed research. LB, BA, AA wrote the manuscript.

## ACKNOWLEDGEMENTS

R00 MH099405 to AA, McNair Scholars Fellowship to AM, Dr. Xuanmao Chen for kindly donating *Gad2-Cre* mice.

